# Extracellular diadenosine tetraphosphate (Ap_4_A) is recognized by the plasma membrane purinoreceptor P2K1/DORN1 and closes stomata in *Arabidopsis thaliana*

**DOI:** 10.1101/2023.05.10.537060

**Authors:** Jędrzej Dobrogojski, Van Hai Nguyen, Joanna Kowalska, Sławomir Borek, Małgorzata Pietrowska-Borek

## Abstract

- Dinucleoside polyphosphates (Np_n_Ns) are considered novel signalling molecules involved in the induction of plant defence mechanisms. However, the Np_n_Ns signal recognition and transduction are still enigmatic. Here we report, for the first time, that diadenosine tetraphosphate (Ap_4_A) is recognized by the *Arabidopsis thaliana* purinoreceptor P2K1/DORN1 (Does Not Respond to Nucleotides 1) and causes stomatal closure.
- Extracellular Ap_4_A- and dicytidine tetraphosphate (Cp_4_C)-induced stomatal closure was observed using a microscope. Reactive oxygen species (ROS) accumulation was determined by staining with nitroblue tetrazolium (NBT) and 3,3ʹ-diaminobenzidine tetrahydrochloride (DAB). Transcriptional changes were determined by quantitative real-time PCR. Wild-type Col-0 and the *dorn1-3 A. thaliana* knockout mutant were used.
- Examination of the leaf epidermis *dorn1-3* mutant provided evidence that P2K1/DORN1 recognizes extracellular Ap_4_A but not Cp_4_C. ROS are involved in signal transduction caused by Ap_4_A and Cp_4_C, leading to stomatal closure. Ap_4_A induced and Cp_4_C suppressed the transcriptional response in wild-type plants. Moreover, in *dorn1-3* leaves, the effect of Ap_4_A on gene expression was impaired.
- Our research demonstrated, for the first time, that P2K1/DORN1 is a plant purinoreceptor for Ap_4_A. This interaction leads to changes in the transcription of signalling hubs in signal transduction pathways.

## Introduction

Regulation of plant metabolic processes takes place at a molecular level. The defence reactions are among the processes in which signal transduction plays a key role. Based on the criterion of the distance that a given signal molecule can cover, short-distance molecules cause local intercellular responses, and long-distance molecules trigger systemic responses. Signalling molecules regulate many processes throughout various signal transduction pathways and specific or unspecific receptors (Sun & Zhang, 2021). Unlike animals, the ability of extracellular nucleotides to initiate diverse signalling responses in plants remained enigmatic for years. A growing number of nucleotides classified as signalling molecules have been identified in plants (Pietrowska-Borek *et al*., 2020a). Among them, extracellular ATP (eATP) plays an essential role in plant growth (Kim *et al*., 2006; Wu *et al*., 2007; Riewe *et al*., 2008; Tonón *et al*., 2010; Clark *et al*., 2010; Zhu *et al*., 2020) and development (Reichler *et al*., 2009; Wu *et al*., 2018). Extracellular ATP regulates responses to biotic stress (Chivasa *et al*., 2005; Chen *et al*., 2017; Tripathi *et al*., 2018; Goodman *et al*., 2022) and abiotic stress (Thomas *et al*., 2000; Kim *et al*., 2009; Sun *et al*., 2012; Hou *et al*., 2018). One of the reactions that eATP can control is stomatal movements (Chen *et al*., 2017; Duong *et al*., 2021; Wang *et al*., 2022). In this reaction, the cytoplasmic Ca^2+^ ions ([Ca^2+^]_cyt_) and the complex signalling cross-talk between second messengers, such as nitric oxide (NO) (Foresi *et al*., 2007; Wu & Wu, 2008; Clark *et al*., 2010), and reactive oxygen species (ROS) (Song *et al*., 2006; Wu *et al*., 2008; Demidchik *et al*., 2009; Chen *et al*., 2017) plays a crucial role, as a mediator in the signal transduction pathway. Consequently, these messenger agents affect the phosphorylation of mitogen-activated protein kinase (MAPK) and the expression of defence-related genes (Choi *et al*., 2014; Chen *et al*., 2017; Li *et al*., 2021).

We have a longstanding interest in the function of dinucleoside polyphosphates (Np_n_Ns) in plant cells. Our papers describe changes in gene expression profile and metabolism in *Arabidopsis thaliana* and *Vitis vinifera* treated with a broad spectrum of Np_n_Ns. We postulated the participation of Np_n_Ns in the plant defence responses since they induce synthesis of the phenylpropanoid pathway-delivered secondary metabolites (Pietrowska-Borek *et al*., 2011, 2014, 2020b). The phenylpropanoid pathway participates in plant defence responses (Dixon & Paiva, 1995; Sharma *et al*., 2019). Identification of Ap_4_A and other Np_n_Ns across prokaryotic and eukaryotic cells testifies to their universality (Ferguson *et al*., 2020). Due to the dramatic increase in levels of various Np_n_Ns observed in cells subjected to abiotic stress factors (Lee *et al*., 1983; Bochner *et al*., 1984; Baltzinger *et al*., 1986; Coste *et al*., 1987; Pálfi *et al*., 1991), these compounds have been termed “alarmones”, triggering stress adaptive processes. Our latest findings confirmed the induction of the phenylpropanoid pathway by purine, pyrimidine, and purine-pyrimidine hybrids of Np_n_Ns. Moreover, we observed that diadenosine polyphosphates (Ap_n_A) induced stilbene biosynthesis. In contrast, dicytidine polyphosphates (Cp_n_C) strongly inhibited this reaction but markedly induced expression of the cinnamoyl-CoA reductase gene that controls lignin biosynthesis (Pietrowska-Borek *et al*., 2020b). Nonetheless, the underlying mechanism of Np_n_Ns signal recognition and transduction in plants remains elusive. The growing number of plant enzymes found to be involved in Np_n_Ns biosynthesis and degradation strengthens the hypothesis of their signalling function (Pietrowska-Borek *et al*., 2020a; Ferguson *et al*., 2020).

Plants can respond to extracellular purine nucleotides, such as eATP, through plasma membrane receptors. So far, two plant receptors with an eATP binding domain have been identified. They are P2K1/DORN1 (Does Not Respond to Nucleotides 1) (Choi *et al*., 2014) and P2K2/DORN2, which belong to the L-type lectin receptor kinase (LecRK) protein family (Pham *et al*., 2020; Cho *et al*., 2023). LecRK proteins activate the processes controlling stress responses, development, growth, and disease resistance (Jose *et al*., 2020). Although eATP sensing and action in plants have been elucidated, the mechanisms of signal perception and transduction evoked by Np_n_Ns, such as Ap_4_A and Cp_4_C, remain enigmatic. In animal cells, among different nucleotides and nucleosides, eATP, together with Ap_4_A, shares access to the same receptors that belong to the P2 group, which is divided into two classes, namely ligand-gated ion channels (P2Xs) and G protein-coupled (P2Ys) receptors (Vigne *et al*., 2000; McDonald *et al*., 2002; Wang *et al*., 2003; Verspohl *et al*., 2010; Burnstock, 2018). Therefore we hypothesise that the purinoreceptor P2K1/DORN1, a receptor of eATP, is also necessary for sensing Ap_4_A in plant cells. Moreover, we wondered whether P2K1/DORN1 can also recognise the pyrimidine nucleotide Cp_4_C.

Here we present, for the first time, evidence for the involvement of the P2K1/DORN1 receptor in the sensing of Ap_4_A in plants. All experiments were conducted on 4-week-old *Arabidopsis thaliana* wild-type Col-0 and *dorn1-3* knockout mutant leaves. Our research showed that extracellular Ap_4_A and Cp_4_C evoked stomatal closure in Col-0 plants. This effect was abolished in the *dorn1-3* mutant by Ap_4_A but not Cp_4_C. This result confirms the requirement of P2K1/DORN1 for Ap_4_A-induced stomatal closure. Nevertheless, our research indicates the involvement of superoxide (^•^O_2_^−^) and hydrogen peroxide (H_2_O_2_) in the signal transduction evoked by Ap_4_A and Cp_4_C, leading to stomatal closure. Furthermore, we analysed the expression of genes encoding selected proteins integrated within the signalling hubs. It concerns NADPH oxidases (*RBOHD* and *RBOHF*), *MAPK* cascades, *SNF1/AMPK*-related protein kinases (*SnRKs*) and transcriptional factors such as *ZAT6* and *ZAT12*. Notably, Ap_4_A induced expression of the tested genes. Moreover, the gene expression in *dorn1-3* was almost abolished by the Ap_4_A effect.

## Materials and Methods

### Nucleotides

Ap_4_A and Cp_4_C were synthesized following previously reported procedures, purified by reversed phase HPLC, and isolated as ammonium (NH_4_^+^) salts. The purities (>95%) were confirmed by analytical HPLC, ^1^H NMR and ^31^P NMR (Pietrowska-Borek *et al*., 2020b).

### Plant material

*Arabidopsis thaliana* lines were in the Columbia (Col-0) ecotype. A T-DNA insertion line of LecRK-I.9 (Salk_042209; *dorn1-3*) was obtained from NASC (Nottingham Arabidopsis Stock Centre, Nottingham, UK). Surface-sterilised seeds were stratified in darkness at 4°C for 48 h and transferred to a growth chamber. Plants were grown for four weeks on the soil at 21-23°C, 60-70% humidity, under a long-day photoperiod (16 h light and 8 h dark), 120 µmol m^-2^ s^-1^ light intensity. Genotyping of insertional mutants is described in Methods **S1**. Primers are listed in Table **S1**.

### Stomatal aperture measurement

To ensure fully open stomata, plants were placed for 3 h under light intensity 120 µmol m^-2^ s^-1^. Samples of leaf epidermis were obtained from the abaxial side. They were placed on a microscope slide for 2 h of incubation in (*i*) MOCK solution MES/KOH opening buffer containing 10 mM MES pH 6.15, 10 mM KCl, 10 μM CaCl_2_ (control), (*ii*) 10 µM abscisic acid (ABA, Sigma Aldrich, A1049) dissolved in the MOCK solution MES/KOH buffer, and (*iii*) 2 mM ADP (Sigma, A2754), ATP (Sigma, AA8937), Ap_3_A, Ap_4_A, and CDP (Sigma, C9755), CTP, Cp_3_C, Cp_4_C dissolved in the MOCK solution MES/KOH buffer. CTP and Np_n_Ns were synthesised as described previously (Pietrowska-Borek *et al*., 2020b). Stomata were observed using the ZOE Fluorescent Cell Imager (Bio-Rad1450031EDU). Measurements, including stomatal aperture width and length, were performed with the ImageJ software. The involvement of ROS in stomatal movement under nucleotide treatment was examined by the simultaneous addition of ROS enzyme scavengers to the nucleotide solutions. Catalase (CAT) (Sigma Aldrich, C100) and superoxide dismutase (SOD) (Sigma Aldrich, S9697), in a concentration of 100 units ml^-1^ and 500 units ml^-1^, respectively, were used together in an incubation mixture.

### Detection of intracellular ROS burst in leaves

Two leaves were incubated in 3 ml of MOCK solution MES/KOH opening buffer or the buffer enriched in 2 mM concentrations of tested nucleotides. After 2 h the incubating buffers were gently replaced with 3 ml of staining solutions and submerged leaves were vacuum infiltrated three times (1 min each time). The staining solution for ^•^O ^−^ detection was composed of 0.5% nitroblue tetrazolium (NBT) dissolved in 10 mM potassium phosphate buffer, pH 7.8 (Müller *et al*., 2009), and the staining solution for H_2_O_2_ synthesis was composed of 3,3′-diaminobenzidine tetrahydrochloride (DAB) (1 mg ml^-1^ DAB) dissolved in 10 mM potassium phosphate buffer, pH 7.4, and 0.05% Tween (Daudi & O’Brien, 2012). Samples were incubated at room temperature for the next 2 h in the dark with continuous shaking. Then, leaves were incubated in 96% ethanol overnight for bleaching, and the photographs were taken with an Epson Perfection V700 scanner.

### Gene expression analyses

According to the manufacturer’s instructions, total RNA was extracted from leaves using the RNeasy Plant Mini Kit (Qiagen). Evaluation of RNA purity, cDNA synthesis, reverse transcription, and RT-qPCR were performed as described previously by Pietrowska-Borek and co-workers (Pietrowska-Borek *et al*., 2011, 2015; Pietrowska-Borek & Nuc, 2013). The qRT-PCR reactions were performed using a CFX96 Real-Time PCR Detection System (Bio-Rad). The specific primers for *Arabidopsis thaliana* genes are listed in Table **S1**. The 2^-ΔΔCt^ method (Schmittgen & Livak, 2008) was applied to calculate the relative gene expression. The data were normalised against the reference gene, *ACTIN2* (*ACT2*). For statistical analysis, the gene expression data were Log_2_-transformed to meet distribution and variance assumptions.

### Statistical analysis

All experiments were performed at least three times. The results are shown as the mean ± SD. The statistical significance of the differences among the means was analysed using Statistica, Version 13 (TIBCO Software Inc., Palo Alto, CA, USA, http://statistica.io).

## Results

### Ap_4_A and Cp_4_C induce stomatal closure

Our previous research showed that exogenous Np_n_Ns induce biosynthesis of secondary metabolites that play an essential role in the plant defence strategy (Pietrowska-Borek *et al*., 2011, 2014, 2020b). We wondered how the signal evoked by Np_n_Ns could be sensed and transduced in plant cells and whether plants contain cell membrane receptor(s) for these molecules. It is known that eATP, one of the exogenous purine nucleotides, evokes stomatal closure with involvement of the purinoreceptor P2K1/DORN1 in *Arabidopsis thaliana* (Choi *et al*., 2014; Chen *et al*., 2017). Therefore based on similarities in ATP and Ap_4_A structure, we tested the effect of these nucleotides on stomatal movements. Moreover, we also included cytosine nucleotides in our research because of the different effects of purine and pyrimidine Np_n_Ns on the phenylpropanoid pathway in *Vitis vinifera* cells (Pietrowska-Borek *et al*., 2020b). To trace stomatal movement under the nucleotide treatment, we examined the ability of purine Np_n_Ns such as Ap_3_A and Ap_4_A to stimulate stomatal closure. Additionally, for the positive control, we tested the effects of ADP and ATP as described earlier (Choi *et al*., 2014; Chen *et al*., 2017) as well as ABA – a well-known molecule controlling stomatal movements (Danquah *et al*., 2014). Exogenous Ap_4_A significantly reduced the stomatal aperture in the light. It was at a similar level compared to the effect evoked by ATP and ADP. However, Ap_3_A did not evoke such an effect (Fig. **1**). We also examined stomatal movement under the treatment of cytidine mono- and dinucleotides (CDP, CTP, Cp_3_C, Cp_4_C). Interestingly, only Cp_4_C triggered significant stomatal closure among tested cytidine nucleotides. As expected, ABA closed stomata (Bharath *et al*., 2021) (Fig. **1**).

**Fig. 1.**
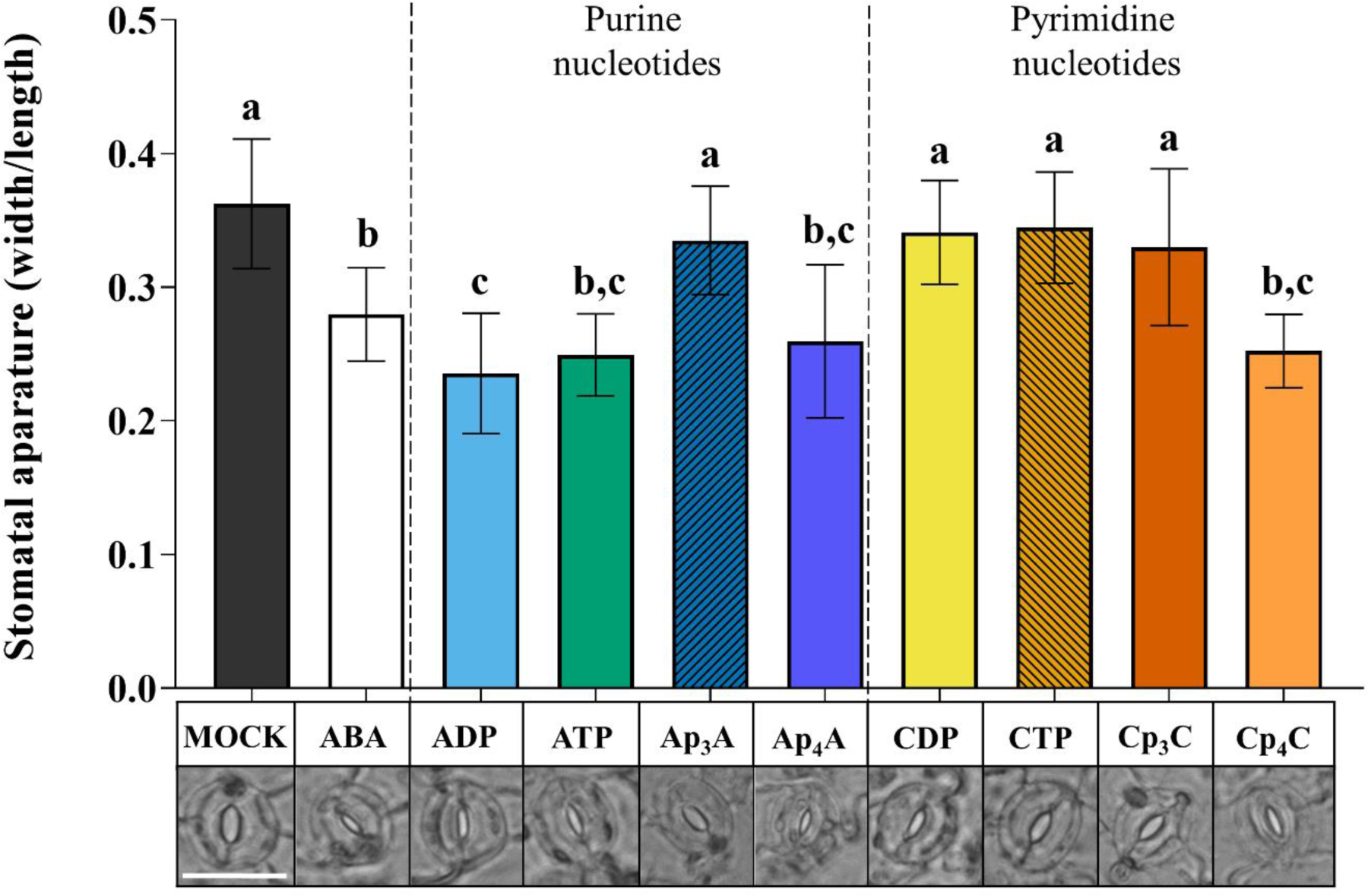
Ap_4_A and Cp_4_C, similarly to ADP and ATP, induce stomatal closure in *Arabidopsis thaliana* Col-0 plants. Images represent stomata in the abaxial epidermis of a leaf treated for 2 h with the MOCK solution MES/KOH opening buffer, 10 µM ABA, and 2 mM purine and pyrimidine nucleotides. White bar = 25 μm. Bars represent mean values ± SD, n ≥ 20, three biological replicates. Different letters above the error bars indicate statistically significant differences according to the ANOVA analysis with the post-hoc Tukey’s HSD multiple comparisons test (p < 0.05).

In plant cells there are enzymes degrading Np_n_Ns to mononucleotides (Guranowski, 2004). To confirm that Ap_4_A and Cp_4_C evoke stomatal closure but not by the products of their degradation (AMP, ADP, ATP, and CMP, CDP, CTP, respectively), we collected samples of leaf epidermis from the microscope slides after incubation of nucleotides, and application of the HPLC assay (Method **S2**) proved that Ap_4_A was not degraded to the corresponding mononucleotide. Only a trace amount of CTP was detected in a solution of Cp_4_C after the investigation (Fig. **S1a,b**).

### P2K1/DORN1 is involved in signal perception evoked by Ap_4_A but not Cp_4_C

Plants respond to eATP by induction of a complex signalling network after signal recognition by the P2K1/DORN1 and P2K2 receptors (Choi *et al*., 2014; Pham *et al*., 2020). Similarities in stomatal movements evoked by eATP, Ap_4_A, and Cp_4_C led us to hypothesise that those nucleotides could interact with P2K1/DORN1. Based on the results presented in Fig. **1**, Ap_4_A and Cp_4_C were chosen for further experiments. The *dorn1-3* mutant, having a T-DNA insertion in the extracellular legume-type lectin domain, was selected based on literature data (Choi *et al*., 2014; Chen *et al*., 2017). We found that Ap_4_A and eATP did not close stomata in *dorn1-3* mutant leaves. Contrary to this, Cp_4_C significantly closed stomata in *dorn1-3* mutant leaves. As expected, ABA-treated mutant leaves also showed closed stomata (Chen *et al*., 2017) (Fig. **2**). Thus, the results strongly suggest that besides eATP, P2K1/DORN1 may also be involved in signal perception elicited by Ap_4_A but not Cp_4_C.

**Fig. 2.**
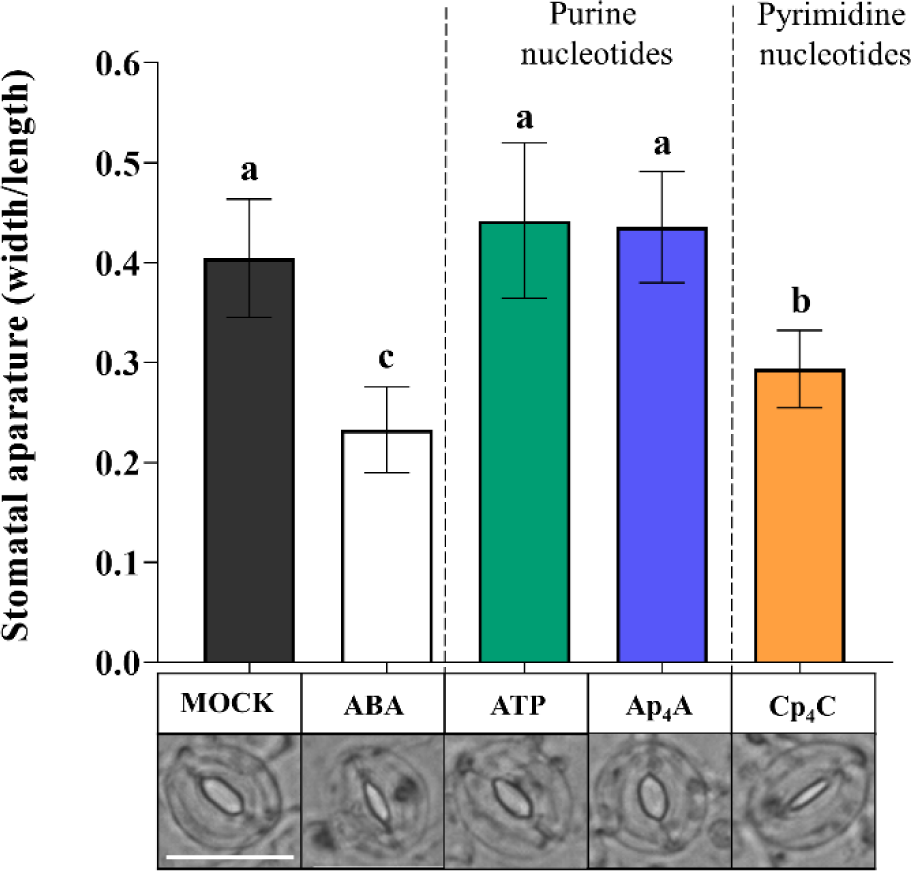
Ap_4_A, similarly to eATP, did not induce stomatal closure in the *dorn1-3 Arabidopsis thaliana* mutant. However, Cp_4_C and ABA evoked stomatal closing. Images represent stomata in the abaxial epidermis of *dorn1-3* leaf treated for 2 h with MOCK solution opening buffer, 10 µM ABA and 2 mM ATP, Ap_4_A, and Cp_4_C. White bar = 25 μm. Bars represent mean values ± SD, n ≥ 20, three biological replicates. Different letters above the error bars indicate statistically significant differences according to the ANOVA analysis with the post-hoc Tukey’s HSD multiple comparisons test (p < 0.05).

### ROS are produced in leaves under nucleotide treatment

It was previously found that elevated production of ROS and stomatal closure are mediated by eATP recognition by the receptor P2K1/DORN1, followed by direct phosphorylation of the NADPH oxidase RBOHD (Chen *et al*., 2017). This phosphorylation causes an increase in generation of extracellular ROS, such as ^•^O_2_^−^, which is then converted into H_2_O_2_ in the extracellular environment (Waszczak *et al*., 2018; Smirnoff & Arnaud, 2019). Notably, the apoplastic production of ROS is one of the fastest physiologically common responses to external stimuli observed in plants (Macho & Zipfel, 2014; Mittler *et al*., 2022). Considering all the above-described information, we decided to investigate the accumulation of ^•^O_2_^−^ and H_2_O_2_ in *Arabidopsis thaliana* leaves in response to 2 mM ATP, CTP, Ap_4_A, and Cp_4_C. Our experiments revealed that blue staining of leaves, indicating ^•^O_2_ ^−^ accumulation, was increased in Col-0 leaves treated with CTP, Ap_4_A, and Cp_4_C but not by eATP, while in the *dorn1-3* mutant, only Cp_4_C evoked accumulation of ^•^O_2_^−^ (Fig. **3a**). Brown staining representing the concentration of H_2_O_2_ in leaves was increased in Col-0 leaves under eATP, Ap_4_A, and Cp_4_C, while CTP caused only slightly brown staining. In the *dorn1-3* mutant, only CTP and Cp_4_C evoked an accumulation of H_2_O_2_ in the leaves. Nevertheless, only weak brown staining was caused by CTP (Fig. **3b**).

**Fig. 3.**
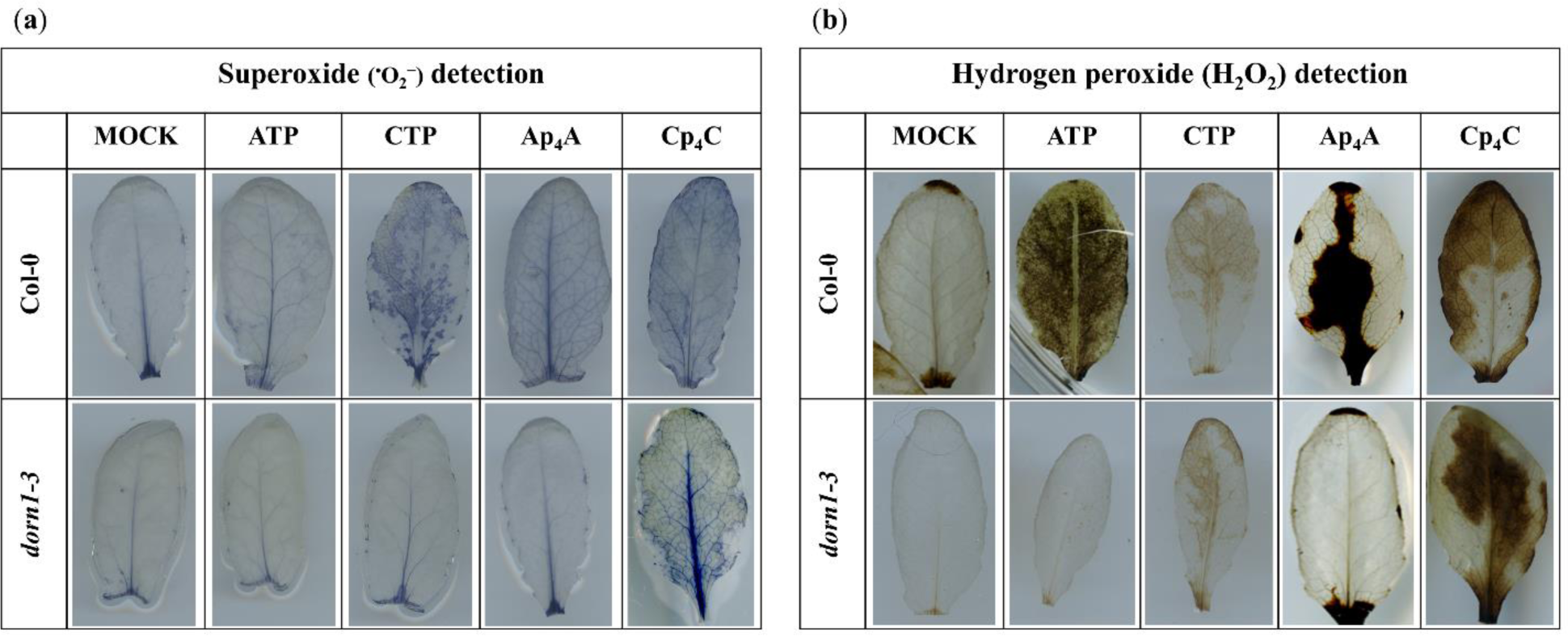
Histochemical detection of (**a**) ^•^O_2_^−^ and (**b**) H_2_O_2_ in leaves of *Arabidopsis thaliana* Col-0 and the *dorn1-3* mutant triggered by 2 mM ATP, CTP, Ap_4_A, and Cp_4_C after 2 h treatment. Leaves were stained with NBT and DAB for ^•^O_2_^−^ and H_2_O_2_ detection, respectively. The experiment was repeated six times, and representative leaves were chosen.

### ROS are involved in signal transduction evoked by eATP, Ap_4_A and Cp_4_C, leading to stomatal closure

Based on the results indicating that Ap_4_A and Cp_4_C induced the production of ROS (Fig. **3a,b**), we wondered whether these key signalling molecules are components of signal transduction pathways evoked by Np_n_Ns leading to stomatal closure. We simultaneously applied superoxide dismutase (SOD) and catalase (CAT), enzymes scavenging ROS (Khokon *et al*., 2011; Mittler *et al*., 2022), and thereby sought to confirm the role of ^•^O ^−^ and H O in the transduction pathway of the signal generated by Ap_4_A and Cp_4_C. Interestingly, CAT and SOD eliminated the effect of stomatal closure under simultaneous nucleotide treatment, so our observations showed direct involvement of ^•^O_2_^−^ and H_2_O_2_ in stomatal closure evoked by eATP, Ap_4_A, and Cp_4_C. However, the plants did close their stomata upon adding ABA (Fig. **4**).

**Fig. 4.**
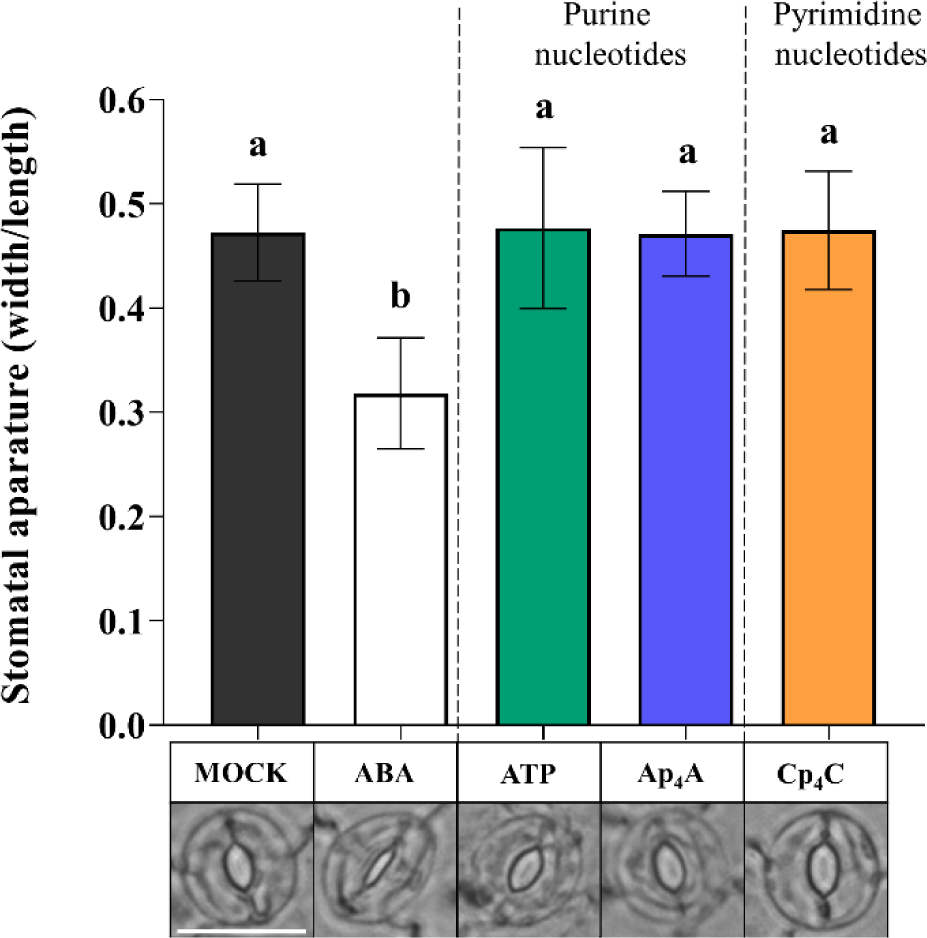
ROS enzyme scavengers, CAT and SOD, eliminate the effect of stomatal closure after the 2 mM ATP, Ap_4_A, and Cp_4_C treatment in *Arabidopsis thaliana* Col-0 leaves. White bar = 25 μm. Bars represent mean values ± SD, n ≥ 20, three biological replicates. Different letters above the error bars indicate statistically significant differences according to the ANOVA analysis with the post-hoc Tukey’s HSD multiple comparisons test (p < 0.05).

### P2K1/DORN1 is implicated in Ap_4_A- and eATP-responsive gene expression

It is known that transcriptional upregulation of defence-related and wound-response genes by eATP is P2K1/DORN1-dependent (Choi *et al*., 2014; Jewell *et al*., 2019). Thus we decided to investigate whether Ap_4_A also changes the expression of the defence-related genes and whether the plasma membrane receptor P2K1/DORN1 is engaged in this regulation. To understand the signal transduction pathway evoked by Ap_4_A, we tested the gene expression coding for proteins as a component of signalling hubs known as key points in response to stresses. First, we studied the NADPH oxidases respiratory burst oxidase homologs (RBOHs), RBOHD, and RBOHF, which generate ROS (Mittler *et al*., 2022). We found that Ap_4_A up-regulated *RBOHF* but not by eATP in Col-0 plants. Interestingly, both eATP and Ap_4_A downregulated *RBOHF* expression in the *dorn1-3* mutant (Fig. **5a**). The expression of *RBOHD* was drastically induced (the most among all studied genes) by eATP but only in Col-0 plants. In contrast, in the *dorn1-3* plants this effect was weak. Ap_4_A evoked slight changes in expression levels of *RBOHD* in Col-0 and *dorn1-3* plants (Fig. **5a**).

**Fig. 5.**
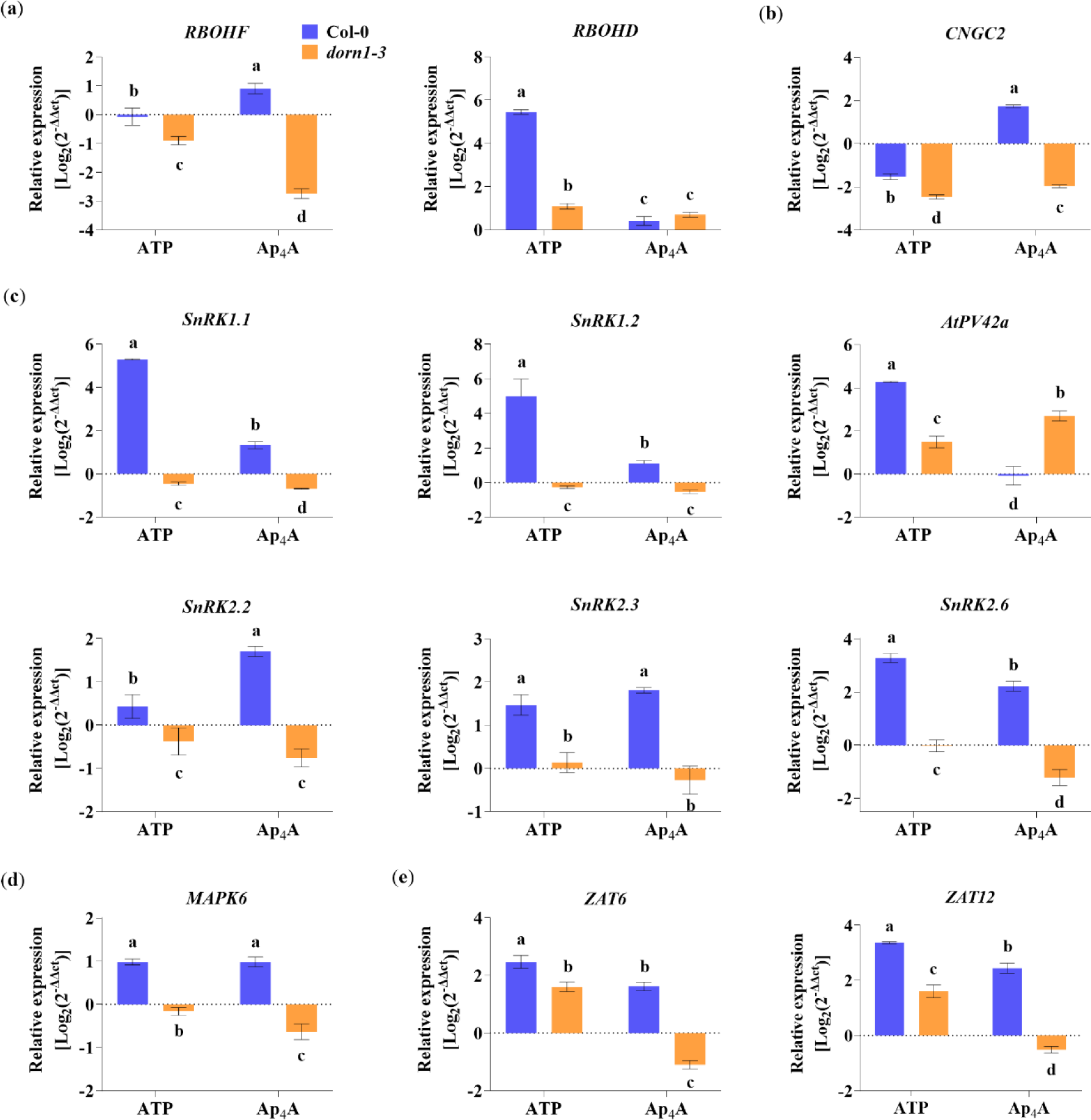
The purinoceptor P2K1/DORN1 is involved in the Ap_4_A-induced transcriptional response in *Arabidopsis thaliana* Col-0 leaves. Graphs present the changes in the gene expression level for (a) NADPH oxidase respiratory burst homologs (*RBOHD* and *RBOHF*), (b) cyclic nucleotide-gated channel 2 (*CNGC2*), (c) SNF1/AMPK-related protein kinases (*SnRKs*), (d) mitogen-activated protein kinase 6 (*MAPK6*), (e) transcription factors (*ZAT6* and *ZAT12*). Leaves taken from Col-0 and the *dorn1-3* mutant were treated for 2 h with 2 mM ATP and Ap_4_A. Transcript levels are represented as Log_2_(2^−ΔΔCt^) compared to the MOCK-treated (control) plants. The housekeeping gene *AtACT2* was used for data normalisation as an endogenous control. Data are mean ± SD from 3 biological replicates. Different letters above the error bars indicate statistically significant differences according to the ANOVA analysis with the post-hoc Tukey’s HSD multiple comparisons test (p < 0.05).

Other components involved in a variety of signalling pathways, ranging from development to stress responses, are cyclic nucleotide-gated channels (CNGCs) (Duszyn *et al*., 2019; Jarratt-Barnham *et al*., 2021). Moreover, AtCNGC2 mediates eATP signal transduction in cells of the root epidermis (Wang *et al*., 2022). We found that Ap_4_A induced *CNGC2* expression in Col-0 plants and decreased the expression in the *dorn1-3* mutant. Extracellular ATP decreased the expression of *CNGC2* in both Col-0 and dorn1-3 mutant plants (Fig. **5b**). We also focused on essential protein kinases, such as SnRKs, that regulate cellular energy homeostasis, stress response, and growth (Zhang *et al*., 2020). Thus, we checked the changes in the expression of *SnRK1.1*, *SnRK1.2*, *SnRK2.1*, *SnRK2.2*, and *SnRK2.6*. We also tested the expression of *PV42a* encoding cystathionine-β-synthase (CBS) domain-containing protein belonging to the PV42 class of γ-type subunits of the plant SnRK1 complexes. It is known that CBS domains generally act as regulatory domains of protein activity through adenosyl ligand binding (Baudry *et al*., 2022). Our experiments showed that eATP strongly induced the expression of *SnRK1.1*, *SnRK1.2*, and *PV42a* in Col-0 plants. Although Ap_4_A causes a lower effect than eATP, the elevation in the expression of *SnRK1.1* and *SnRK1.2* was statistically significant. Interestingly, in Col-0 plants, only eATP up-regulates the transcription of *PV42a*. Still, in the *dorn1-3* mutant compared to Col-0, only Ap_4_A treatment caused induction of the expression (Fig. **5c**). Extracellular ATP and Ap_4_A increased the expression of *SnRK2.2*, *SnRK2.3*, and *SnRK2.6* in Col-0 plants. In the *dorn1-3* mutant plants, Ap_4_A down-regulated *SnRK2.2*, *SnRK2.3*, and *SnRK2.6*. Still, the effect of eATP in the mutant was not the same for the expression of the three *SnRK2* genes; namely, the expression of *SnRK2.2* was decreased, *SnRK2.3* was slightly increased, and there was no effect on *SnRK2.6* expression (Fig. **5c**). The strong relationships between secondary messengers, such as ROS and MAPKs, are often highlighted in the literature (Matsushita *et al*., 2020; Byrne *et al*., 2020). MAPK6, among its roles in various metabolic processes in plants, can regulate the activities of diverse targets, including transcription factors (Smékalová *et al*., 2014). We observed up-regulation of *MAPK6* expression by both eATP and Ap_4_A in Col-0 plants and down-regulation in the *dorn1-3* mutant (Fig. **5d**). Among transcription factors that MAPKs regulate, we tested the regulation of expression of the zinc-finger transcription factors (*ZAT6* and *ZAT12*) and we found that eATP and Ap_4_A up-regulated the expression of both genes, as mentioned above in Col-0 plants. Extracellular ATP increased the expression of *ZAT6* and *ZAT12* also in the *dorn1-3* mutant, but Ap_4_A downregulated the expression of both genes in the mutant plants (Fig. **5e**).

## Discussion

Plants are exposed to continuous changes in environmental conditions that lead to an imbalance in cellular homeostasis. It is known that in response to various stresses in prokaryotic and eukaryotic cells, Np_n_Ns accumulate. The accumulation of such uncommon nucleotides can be considered in the context of the “friend hypothesis” (alarmone) and “foe hypothesis” regarding critically damaged cells as a result of internal and external stresses (McLennan, 2000; Ferguson *et al*., 2020). Although there are identified Ap_4_A-binding protein targets in cells (Ferguson *et al*., 2020), the signalling pathways are still unclear. We reported previously that extracellular Np_n_Ns regulate the phenylpropanoid pathway producing secondary metabolites – key molecules in response to abiotic stress in *Arabidopsis thaliana* and *Vitis vinifera* (Pietrowska-Borek *et al*., 2011, 2014, 2020b). Notably, one of the phenylpropanoid pathway enzymes, 4-coumarate:CoA ligase, is known to catalyse synthesis of Ap_4_A (Pietrowska-Borek *et al*., 2003), and its activity was increased by Ap_4_A (Pietrowska-Borek *et al*., 2011). It is known that some extracellular Ap_n_N may become internalised and operate intracellularly (Ferguson *et al*., 2020). Despite this obvious evidence of the signalling function of uncommon nucleotides in regulating phenylpropanoid synthesis, no receptors or signalling pathways have been identified in plants until now. Here we demonstrated, for the first time, that Ap_4_A evoked stomatal closure in *Arabidopsis thaliana* leaves (Fig. **1**). We did not observe such an effect in *dorn1-3* plants (Fig. **2**). So we can conclude that plasma membrane purinoreceptor P2K1/DORN1 recognises Ap_4_A. However, our research also indicates that P2K1/DORN1 is not involved in signal perception elicited by Cp_4_C (Fig. **2**). Such results suggest that in plants, there is not or is other Cp_4_C-binding protein(s). After Ap_4_A signal recognition, P2K1/DORN1 stimulates ROS burst and the defence-related response. Our data indicating ROS involvement in the plant response to Ap_4_A and Cp_4_C support the hypothesis concerning the signalling function of Np_4_Ns (Fig. **3** and Fig. **4**). Moreover, the HPLC assay proved that Ap_4_A was not degraded to corresponding mononucleotides, which could evoke stomatal closure during the experiment (Fig. **S1**). Only a tiny amount of CTP was detected in a solution of Cp_4_C after the investigation. Still, as we proved, CTP did not evoke stomatal closure (Fig. **1**). Therefore, it confirms that the observed stomatal closure and ROS accumulation were caused by Ap_4_A and Cp_4_C but not by their decomposition products.

The upregulation of defence-related genes encoding proteins involved in signalling hubs was reported (Zhang *et al*., 2020). The expression of the genes described in this research was mostly abolished or down-regulated in the *dorn1-3* mutant (Fig. **5**). Recent studies consider cross-talk between diverse plant defence response markers such as ROS, hormones, and kinase cascades, leading to transcriptional, translational, and metabolic reprogramming (Mittler *et al*., 2022). Our transcriptional analysis focused on elements that integrate various signals and included cyclic nucleotide-gated channels (CNGC) and NADPH oxidases – respiratory burst oxidase homologs (RBOHD, and RBOHF) that generate ROS. Moreover, our studies are focused on SNF1-related protein kinases (SnRKs) and PV42a, a cystathionine-β-synthase (CBS) domain-containing protein belongs to the PV42 class of γ-type subunits of the plant SnRK1 complexes. The next elements of signal transduction pathways that we tested concern MAPK6, and transcription factors, ZATs (Fig. **5**). The transcript level of *CGNC2* increased only under Ap_4_A in Col-0 plant leaves (Fig. **5b**). Involving CGNC2 in another purine nucleotide, eATP, signal transduction in the root epidermis and eATP-induced Ca^2+^ influx were described by Wang (Wang *et al*., 2022). This result suggests that CNGC channels can be a part of signal transduction evoked by Ap_4_A.

Rapid systemic signalling in response to stress can be stimulated by RBOHD and RBOHF, producing apoplastic ROS (Choi *et al*., 2016). It is known that the elevated production of ROS and stomatal closure are mediated by eATP recognition by the receptor P2K1/DORN1, followed by direct phosphorylation of RBOHD (Chen *et al*., 2017), while *RBOHD* expression was significantly reduced in *dorn1-3* mutant plants. Our studies showed that transcriptomic changes in both *RBOHD* and *RBOHF* evoked by Ap_4_A are similar, but in the *dorn1-3* plants, expression of *RBOHF* also was strongly inhibited (Fig. **5a**). This observation correlated with the accumulation of ROS in *Arabidopsis thaliana* leaves (Fig. **3**). Stress signalling in plants also involves different families of kinases, including the MAPK module, that can be activated by ROS (Meng & Zhang, 2013). Moreover, it was previously shown that MAPKs are activated by eATP (Choi *et al*., 2014; Medina-Castellanos *et al*., 2014; Chen *et al*., 2021; Cho *et al*., 2022). We observed the induction of *MAPK6* expression evoked by eATP and Ap_4_A (Fig. **5d**), and it is known that MPK6 modulates actin remodelling to activate stomatal defence in *Arabidopsis thaliana* (Zou *et al*., 2021). MAPK pathways are necessary for several ABA responses in many plant species, including antioxidant defence and guard cell signalling (Danquah *et al*., 2014). A complex protein complex, SNF1-related protein kinase 1s (SnRK1s) and SnRK2s, plays a prominent role in ABA signalling (Jossier *et al*., 2009; Ou *et al*., 2022). Numerous studies indicate SnRK1s and SnRK2s as regulators of the target of rapamycin (TOR) kinase activity in controlling autophagy (Signorelli *et al*., 2019; Belda-Palazón *et al*., 2020). We observed that Ap_4_A induced the expression of both *SnRK1*s and *SnRK2*s at a similar level in Col-0 plants. However, induction evoked by eATP was much higher for *SnRK1*s than *SnRK2*s in wild-type plants. In the *dorn1-3* mutant, the expression of *SnRK1*s and *SnRK2*s was decreased (Fig. **5c**). Also both tested pyrimidine nucleotides, CTP and Cp_4_C, did not affect expression of *SnRK*s in Col-0 plants (Fig. **S2**). It is known that SnRKs can regulate RBOH, which is engaged in ROS production (Mittler *et al*., 2022). The SnRK1s and SnRK2s were identified as critical nodes for stress and growth signalling pathways (Zhang *et al*., 2020). Moreover, it was suggested that under normal conditions, cytosol-localised SnRK1.1, in response to high-ammonium or low-pH stress, migrates to the nucleus and promotes the phosphorylation of the transcription factors regulating the expression of responsive genes (Sun *et al*., 2021). Studies on AKINβ1, subunit SnRK1, showed its regulatory effect on secondary metabolic processes (e.g. phospholipid and flavonoid metabolism) (Wang *et al*., 2020). Another SnRK1 subunit is PV42a, which is the CBS domain protein. Ap_4_A did not change the expression of the gene encoding AtPV42a in Col-0 plants (Fig. **5c**). It is known that enzymes containing CBS domains can be regulated by Ap_4_A binding (Ferguson *et al*., 2020). Therefore we postulate that AtPV42a regulates SnRK1s in response to Ap_4_A. Moreover, SnRK1, SnRK2, and MAPK interact with transcriptional factors (Zhang *et al*., 2022; Son *et al*., 2023). The induction of ZAT12 and ZAT6 transcription factors in which MAPK6 is involved in an abiotic stress marker was described (Smékalová *et al*., 2014). In the present research, we found that Ap_4_A and eATP induced both *ZAT6* and *ZAT12* gene expression in Col-0 plants, and lack of the P2K1/DORN1 receptor in the *dorn1-3* mutants diminished this effect (Fig. **5e**). It is known that the transcript level of *ZAT6* positively affected the concentrations of phenylpropanoids, including anthocyanin and total flavonoids (Shi *et al*., 2018). Moreover, it was proved that ZAT6 and ZAT12 are involved in the response to cadmium stress and abiotic stress in plants (Opdenakker *et al*., 2012; Shi *et al*., 2014; Chen *et al*., 2016; Dang *et al*., 2022) and the expression of *ZAT12* was strictly dependent on the ROS wave (Brumbarova *et al*., 2016; Myers *et al*., 2022).

The results of our research presented here shed more light on the signalling function of Ap_4_A, its perception and signal transduction pathway in plants. We had previously proposed a hypothetical Np_n_Ns signalling network in a plant cell. Then we strongly suggested the existence of some receptor and signalling transduction pathways involving signalling hubs and transcription factors resulting in gene expression changes, including genes coding for enzymes catalysing the phenylpropanoid pathway (Pietrowska-Borek *et al*., 2011, 2014, 2020b,a). Here, we fill a few gaps in this network (Fig. **6**). Nevertheless, further studies are required to fully describe the role of Np_n_Ns in signalling hubs and better understand the function of uncommon nucleotides in plants.

**Fig. 6.**
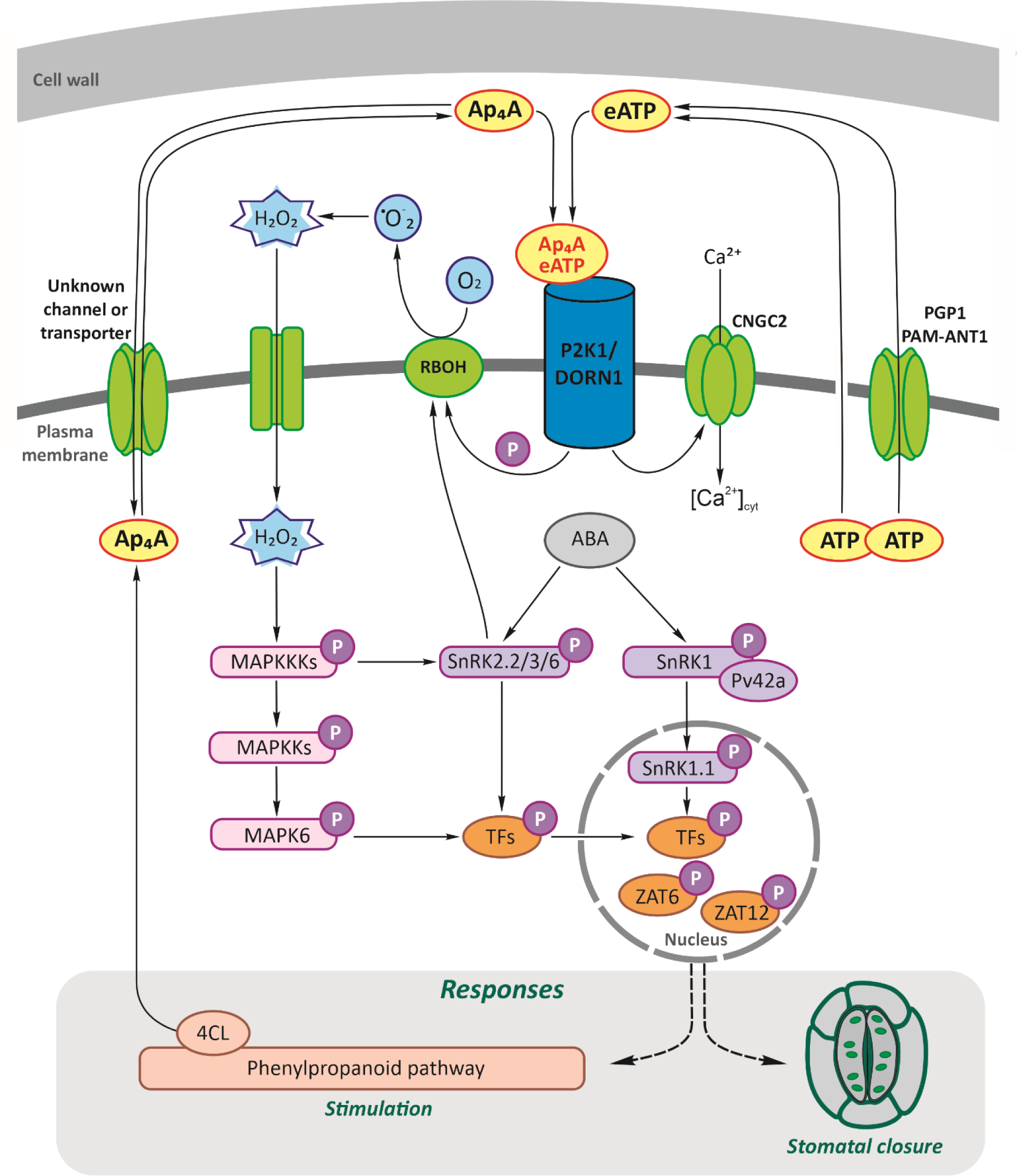
Hypothetical working model of Ap_4_A signalling network in a plant cell. Ap_4_A, similarly to eATP (Choi *et al*., 2014), can be recognized by the purinoreceptor P2K1/DORN1, and lead to stomatal closure. As our study showed, Ap_4_A triggered the ROS wave, which evoked changes in the expression of the defence-related genes encoding proteins involved in signalling hubs, such as CNGC2; RBOHD and RBOHF generate ROS; SnRKs; AtPV42a, γ-type subunits of the plant SnRK1 complexes; MAPK cascades, and transcription factors, ZATs. The wounded cell membrane and transporters can release ATP to the extracellular space matrix: PGP1, p-glycoprotein belonging to ATP-binding cassette ABC transporters, and PM-ANT1, plasma membrane-localized nucleotide transporters (Thomas *et al*., 2000; Rieder & Neuhaus, 2011). Extracellular ATP recognition by P2K1/DORN1 evoked phosphorylation of RBOHD (Chen *et al*., 2017). Also, CNGC2 (Wang *et al*., 2022) and MAPK cascades are involved in eATP signal transduction (Choi *et al*., 2014; Medina-Castellanos *et al*., 2014; Chen *et al*., 2021; Cho *et al*., 2022). We previously described that 4-coumarate:CoA ligase (4CL), the branch point of the phenylpropanoid pathway, can synthesize Ap_4_A (Pietrowska-Borek *et al*., 2003), and its activity is induced by Ap_4_A (Pietrowska-Borek *et al*., 2011). As yet, no channel or transporter for Ap_4_A in plants is known. PM, plasma membrane; P, phosphate.

## Acknowledgements

The publication was co-financed within the framework of the Polish Ministry of Science and Higher Education’s programme “Regional Excellence Initiative”, in the years 2019–2023 (No. 005/RID/2018/19)”, financing amounting PLN 12 000 000 to MPB, and the programme “Inicjatywa doskonałości – Uczelnia Badawcza” granted to Adam Mickiewicz University Poznań, Poland to SB. VHN is grateful for the funding from the European Union’s Horizon 2020 research and innovation programme under the Marie Skłodowska-Curie grant agreement No 861381 (NATURE-ETN).

## Author contributions

JD co-designed the studies, carried out experiments, analysed results, and was involved in statistical analysis, visualisation, and writing of the original draft of the manuscript; VHN and JK synthesized the dinucleoside polyphosphates and participated in reviewing and editing of the manuscript; SB participated in writing and critically reviewing the manuscript, and co-designed and prepared Fig. 6; MP-B conceived the topic of the research, planned and supervised all experiments, analysed all results, performed the statistical analysis, participated in writing the draft of the manuscript, co-designed and co-created all figures, and prepared the final version of the manuscript. All authors read and approved the manuscript.

## Acknowledgement

This work was partially supported by the statutory activity of Poznań University of Life Sciences, no. 506.181.01 to M.P.-B. and J.D. and no. 506.181.09 to M.P.-B.

We thank Richard Ashcroft (bioscience editor, www.anglopolonia.com/home.html) for the professional language editing of the manuscript.

## Data availability

The original data obtained during this research are available from the corresponding authors on reasonable request, and some of them are also accessible in the Supplementary materials.

## Support Information

Additional Supporting Information my be found online in the Supporting Information section at the end of the article.

**Fig. S1** High-performance liquid chromatography analysis of Ap_4_A and Cp_4_C solutions after 2 h of leaf epidermal peel treatment.

**Fig. S2** CTP and Cp_4_C do not up-regulate expression of *SnRK* genes.

**Methods S1** Genotyping *dorn1-3* insertional mutant.

**Method S2** High-performance liquid chromatography (HPLC) analysis of 2 mM solutions of Ap_4_A and Cp_4_C after 2 h leaf epidermal peel treatment.

**Table S1** List of primers used for genotyping T-DNA mutant and qPCR.

Please note: Wiley Blackwell are not responsible for the content or functionality of any Supporting Information supplied by the authors. Any queries (other than missing material) should be directed to the New Phytologist Central Office.

