## Supplementary material for "Extracellular diadenosine tetraphosphate (Ap_4_A) is recognized by the plasma membrane purinoreceptor P2K1/DORN1 and closes stomata in *Arabidopsis thaliana*": Supporting Information_New_Phytologist_MPB.pdf

**New Phytologist Supporting Information**

Article acceptance date:

**Methods S1 Genotyping *dorn1-3* insertional mutant.**

A T-DNA insertion line of LecRK-I.9 (Salk_042209; *dorn1-3*) was obtained from the Nottingham Arabidopsis Stock Centre (NASC, UK). Genomic DNA was extracted from 4-week-old *Arabidopsis thaliana* leaves using a slightly modified hexadecyltrimethylammonium bromide (CTAB) protocol (Doyle & Doyle, 1987). Homozygosity for the T-DNA insertion was confirmed by PCR-based genotyping using the specific primers listed in Table **S1**. PCR conditions were as follows: 35 cycles at 95°C for 30 s, 50°C for 15 s, and 72°C for 1:40 min. Then, gel electrophoresis (1% (w/v) agarose in 1X TBE buffer) was conducted for gene product visualization and final homozygosity confirmation.

**Table S1 List of primers used for genotyping T-DNA mutant and qPCR.**

| Purpose | Gene ID (TAIR) | Primers | Sequence (5′→3′) | Product length | Source |
|---|---|---|---|---|---|
| **Genotype** | | | | | |
| *p2k1-3* mutant | **AT5G60300** | *mlecat5g60300-s* | TCCATGCAACAGTTGCGTTGTCT | | (Choi *et al.*, 2014) |
| | | *P2K1_R* | CTGCAATACCCAAACAGTGGTA | | |
| | | *LBb1* | GCGTGGACCGCTTGCTGCAACT | | |
| **qPCR** | | | | | |
| | **AT3G01090** | *SnRK1.1_F* | CCGCTCCAGAGGTAATTTCG | 68 bp | (Sun *et al.*, 2021) |
| | | *SnRK1.1_R* | CACACCACAGCTCCAGACATCT | | |
| | **AT3G29160** | *SnRK1.2_F* | CACCATTCCTGAGATCCGTCA | 66 pb | |
| | | *SnRK1.2_R* | GAGACAGCAAGATAACGAGGGAG | | |
| | **AT3G50500** | *SnRK2.2_F* | ATATGCCATCGGGATCTGAA | 115 bp | (Nakashima *et al.*, 2009) |
| | | *SnRK2.2_R* | TTGGTTGGGAATGAAGAACAG | | |
| | **AT5G66880** | *SnRK2.3_F* | GTTGGATGGAAGTCCTGCTC | 146 bp | |
| | | *SnRK2.3_R* | TGCCATCATATTCCTGACGA | | |
| | **AT4G33950** | *SnRK2.6_F* | CACAGGAAGCTTGGACATAGAT | 94 pb | (Ou *et al.*, 2022) |
| | | *SnRK2.6_R* | GTACACAATCTCTCCGCTACTG | | |
| | **AT1G15330** | *AtPV42a_F* | GGGATTCTCACGATGCTTGAC | 135 bp | (Fang *et al.*, 2011) |
| | | *AtPV42a_R* | TGTCCAGAGACTGAGTCCTTCG | | |
| | **AT2G43790** | *MAPK6_F* | ACGATGCCATAAGCACCCTTGC | 161 bp | (Kannan *et al.*, 2012) |
| | | *MAPK6_R* | GCGGCTCCATCGCCTCAGAT | | |
| | **AT5G04340** | *ZAT6_F* | AAACCGTGACCTTGACCTGC | 300 bp | (Shi *et al.*, 2018) |
| | | *ZAT6_R* | CTCCGTTCTTTCCTTCGTAGTG | | |
| | **AT5G59820** | *ZAT12_F* | GAGTCACAAGAAGCCTAACAACGA | 242 bp | (Le *et al.*, 2016) |
| | | *ZAT12_R* | AAGCCACTCTCTTCCCACTGCTA | | |
| | **AT5G15410** | *CNGC2_F* | TCTTCAGGTGGATTGGACTGT | 89 bp | (Wang *et al.*, 2022) |
| | | *CNGC2_R* | TCCACCGTTGATTTGGAGGT | | |
| | **AT5G47910** | *RBOHD_F* | CATGCGGGTGCCCATTT | 51 bp | (Choi *et al.*, 2014) |
| | | *RBOHD_R* | ATCCGCGGCAATTAAACG | | |
| | **AT1G64060** | *RBOHF_F* | CTTGGCATTGGTGCAACTCC | 151 bp | (Morales *et al.*, 2016) |
| | | *RBOHF_R* | TCTTTCGTCTTGGCGTGTCA | | |
| | **AT3G18780** | *ACT2_F* | ACTTTCATCAGCCGTTTTGA | 190 bp | (Pietrowska-Borek *et al.*, 2015) |
| | | *ACT2_R* | ACGATTGGTTGAATATCATCAG | | |

**Method S2 High-performance liquid chromatography (HPLC) analysis of 2 mM solutions of $Ap_4A$ and $Cp_4C$ after 2 h leaf epidermal peel treatment.**

Potential degradation of $Ap_4A$ and $Cp_4C$ after 2 h incubation on a slide with a leaf was tested. Samples were analysed for purine nucleotides according to (Guranowski *et al.*, 2009) with minor modifications. Samples of $Ap_4A$ solution (20 µl) were diluted three times with $K_2/KH_2PO_4$ buffer and filtered (Anopore 0.2 µm), and analysed by HPLC in the UV–VIS range using a Discovery C18 column (4.6 x 250 mm, 5 µm; Supelco); flow rate 1 $ml \cdot min^{-1}$. Gradient elution was performed with 0.1 M $KH_2PO_4$, pH 6.0 (solvent A); solvent A/methanol (9: 1, v/v) (solvent B): 0–9 min, 0% B; 9–15 min, 25% B; 15–17.5 min, 90% B; 17.5–19 min, 100% B; 19–23 min, 100% B and 23–30 min, 0% B. Solutions of pyrimidine nucleotides, $Cp_4C$, were analysed according to (Guranowski *et al.*, 2008) with minor modifications. The samples were chilled, diluted three times with 50 mM TEAB (triethylamine buffer, pH 7.4), and filtered (Anopore 0.2 µm). Then, samples were analysed by HPLC in the UV–VIS range using a Discovery C18 column (4.6 x 250 mm, 5 µm; Supelco); flow rate 1 $ml \cdot min^{-1}$. The column was eluted with a linear gradient of 50 mM TEAB (pH 7.4) (solvent A) and solvent A:acetonitrile (60:40, v/v) (solvent B); 0–19 min, 40% B. Nucleotides were identified (purines at 260 nm and pyrimidines at 271 nm) and quantified by comparison with respective standards.

**Fig. S1 High-performance liquid chromatography analysis of $Ap_4A$ and $Cp_4C$ solutions after 2 h of leaf epidermal peel treatment.**

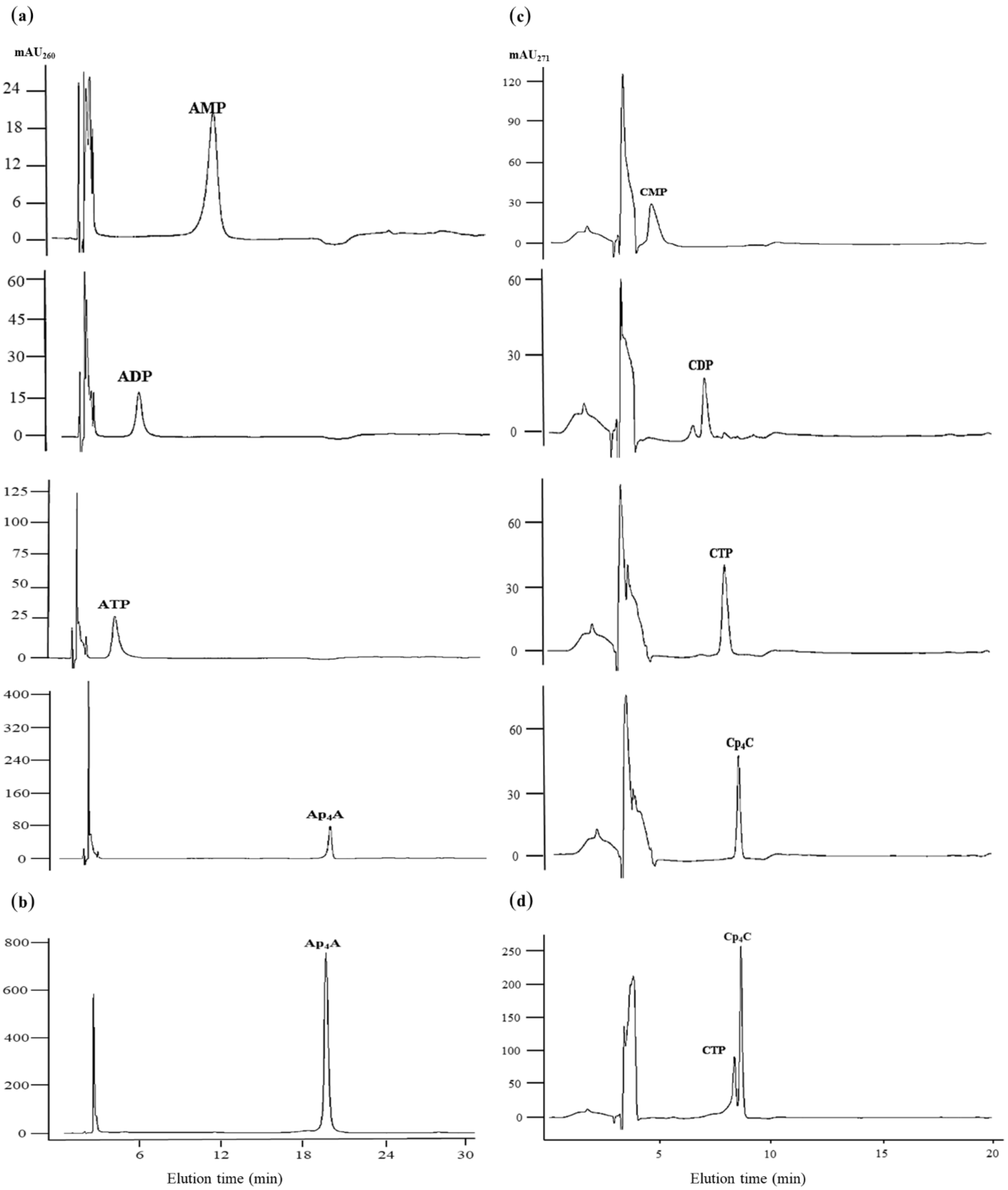

High-performance liquid chromatography (HPLC) analysis was carried out to elucidate Ap$_4$A and Cp$_4$C potential degradation during 2 h of treatment. Samples were analysed for purine nucleotides according to (Guranowski *et al.*, 2009), and samples were analysed for pyrimidine nucleotides according to (Guranowski *et al.*, 2008). (**a**) Chromatography of adenine (AMP, ADP, ATP and Ap$_4$A) and (**c**) cytidine (CMP, CDP, CTP and Cp$_4$C) nucleotide standards, and (**b**) Ap$_4$A and (**d**) Cp$_4$C solution from microscope slide after 2 h of epidermal peel treatment. As we observed, a small amount of CTP was detected only in the sample containing Cp$_4$C.

**Fig. S2 CTP and Cp$_4$C do not up-regulate expression of *SnRK* genes.**

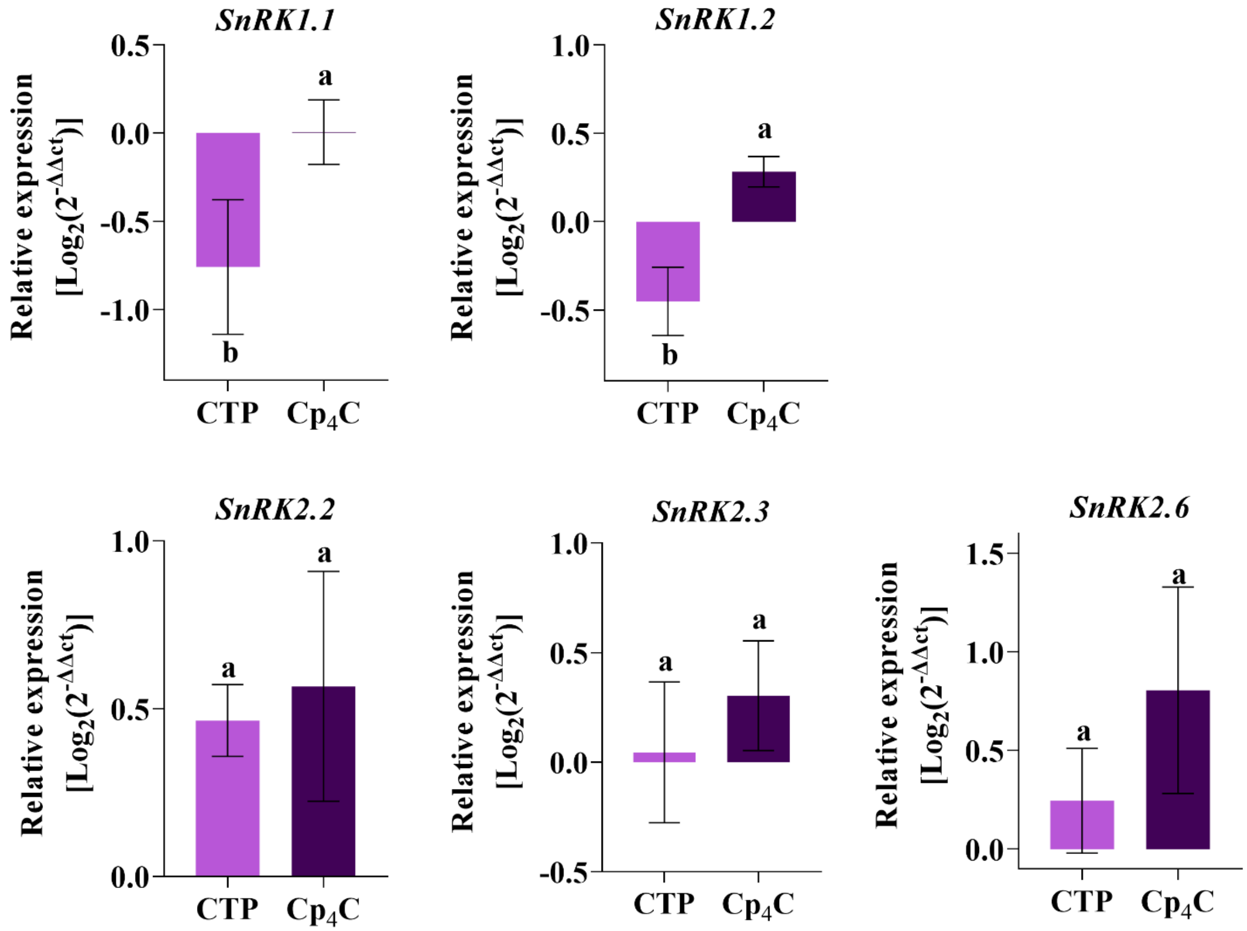

Relative gene expression in 4-week-old leaves of Col-0 and *dorn1*-3 treated with a 2 mM solution of CTP and $Cp_4C$ for 2 h. Afterwards, the total RNA was isolated from leaves and transcribed into cDNA, which was used as a template for quantitative real-time PCR, according to the description in the Material and Methods. Transcript expression is represented as the $\log_2(2^{-\Delta\Delta Ct})$ compared to MOCK-treated plants. The housekeeping gene *AtACT2* was used for data normalisation as an endogenous control. Data are mean ± SD from three independent trials within >3 biological replicates. According to the ANOVA statistical analysis and Tukey's HSD multiple range test ($p < 0.05$), values without a common superscript are statistically significant.
